# DiffuST: a latent diffusion model for spatial transcriptomics denoising

**DOI:** 10.1101/2024.06.19.599672

**Authors:** Shaoqing Jiao, Dazhi Lu, Xi Zeng, Tao Wang, Yongtian Wang, Yunwei Dong, Jiajie Peng

## Abstract

Spatial transcriptomics technologies have enabled comprehensive measurements of gene expression profiles while retaining spatial information and matched pathology images. However, noise resulting from low RNA capture efficiency and experimental steps needed to keep spatial information may corrupt the biological signals and obstruct analyses. Here, we develop a latent diffusion model DiffuST to denoise spatial transcriptomics. DiffuST employs a graph autoencoder and a pre-trained model to extract different scale features from spatial information and pathology images. Then, a latent diffusion model is leveraged to map different scales of features to the same space for denoising. The evaluation based on various spatial transcriptomics datasets showed the superiority of DiffuST over existing denoising methods. Furthermore, the results demonstrated that DiffuST can enhance downstream analysis of spatial transcriptomics and yield significant biological insights.

## 1 Background

Spatial transcriptomics technologies can obtain spatial context and transcriptional pattern of cells in a tissue simultaneously [1, 2], which have become powerful tools for understanding biological mechanisms [3] and disease pathology [4]. Currently, various techniques have been developed for spatial transcriptomics, including 10X Visium [5], Slide-seq [6], stereo-seq [7], and light-seq [8]. For most techniques, in addition to gene expression data and location information, the matched pathology images stained by hematoxylin and eosin (H&E) or immunofluorescence (IF) are also provided. These technologies have made it possible to uncover the complex transcriptional architecture within tissues, characterize spatial expression patterns, and enhance our understanding of cellular mechanisms in diseases [9, 10]. Unfortunately, noise resulting from low RNA capture efficiency [11] and experimental steps needed to keep spatial information [12] may corrupt the biological signals and obstruct analyses. Consequently, denoising becomes a necessary step before the downstream analysis of spatial transcriptomics data.

Most spatial transcriptomics technologies yield three interrelated yet distinct categories of data, including imaging data, gene expression profiles, and data on spatial positioning of gene expression profiles. Among the three types of data, noise is mainly involved in the gene expression profiles. Currently, most denoising methods for spatial transcriptomics data focus on gene expression profiles. Existing methods for denoising spatial transcriptomics data can be broadly categorized into two types. One type removes the noise of spatial transcriptomics data relying on the paired single-cell RNA-seq (scRNA-seq) data [13–17]. For example, Seurat [13] applies canonical correlation analysis [18] to embed spatial transcriptomics and scRNA-seq data into a common latent space, projecting cells from scRNA-seq to the spots of the spatial transcriptomics to reconstruct spatial expression patterns. Tangram [14] uses non-convex optimization and a deep learning framework to learn a spatial alignment of scRNA-seq data. STEM [17] employs a deep transfer learning model to generate spatially-aware embedding of both scRNA-seq and spatial transcriptomics data, facilitating precise spatial expression patterns denoising. This type of method requires the matched scRNA-seq data from the same tissue, which needs extra experimental cost and is not always available. The other type utilizes the location information and imaging data to enhance the gene expression profiling [11, 12, 19–21]. For instance, GraphST [19] combines graph neural networks with self-supervised contrastive learning to generate discriminative spot representations and achieve denoising by minimizing the embedding distance between spatially adjacent spots. MIST [20] combines molecular similarity and spatial connectivity between spots to denoise spatial transcriptomics data through a region-based low-rank approximation approach. Sprod [12] utilizes positional and image features as input, constructs a latent graph from these features, and subsequently employs the latent graph to smooth out noise in the original spatial transcriptomics expression matrix. However, most of these methods primarily focus on positional information, ignoring the rich structural and semantic information involved in the imaging data.

For a given tissue, the imaging data and gene expression profiles are different views from different scales. Therefore, we hypothesize that using the rich information included in imaging data can enhance the gene expression profiles. Here, the key challenge is mapping these two data types with different scales to the same space for denoising. To address this challenge, we developed a latent diffusion model for spatial transcriptomics denoising, named DiffuST. The model is to systematically and slowly destroy the structure in spatial transcriptomics data distribution through an iterative forward diffusion process [22]. At each time step, Gaussian noises are added to the data. The denoising process predicted the noise in each diffusion step to denoise spatial transcriptomics data through a learned reverse process [23, 24]. Furthermore, the model leverages the characteristics of latent diffusion models to integrate different types of features into the same space, resolving the semantic inconsistencies arising from feature fusion in high-dimensional space [25–27]. Moreover, due to the disparate scales of positional information and imaging data, DiffuST employs different strategies for feature extraction. Specifically, unlike traditional models of directly using pixel intensity or texture features as image features, DiffuST utilizes a pre-trained model to extract enriched high-level semantic features. For positional information, DiffuST utilizes a graph autoencoder to capture the spatial relations among the spots. By testing on various spatial transcriptomics datasets, we showed its superiority over existing denoising methods. The results demonstrated its superiorities for downstream analysis tasks such as tissue structure identification, inference of cell-to-cell communications, and deconvolution of cell types. Overall, DiffuST is a powerful method for spatial transcriptomics denoising, which enables enhanced downstream analysis and provides valuable insights into complex biological systems.

## 2 Results

### 2.1 Overview of DiffuST

In this section, we provide a brief overview of DiffuST, a latent diffusion model designed to integrate spatial information and image features for spatial transcriptomics denoising (Fig. 1). DiffuST comprises three major modules: spatial information compression and gene expression profiles reconstruction module (Fig. 1a); latent representation denoising module (Fig. 1b); image features conditioning mechanism module (Fig. 1c).

Firstly, we leverage the spatial information of the spatial transcriptomics dataset to construct an undirected neighborhood graph where spots close to each other are connected. Next, the encoder of a graph autoencoder is built to effectively compress the gene expression profiles and spatial similarity into a latent representation *z* by iteratively aggregating gene expressions from neighboring spots (Fig. 1a). Following this, a latent diffusion model is designed to generate a denoising latent representation *ẑ*. During the training process, starting with latent representation *z*, Gaussian noise is iteratively added over *T* discrete steps until *z_T_* . This noising procedure is performed via the Markov process *q*(*z_t_|z_t−_*_1_). The diffusion model is trained to predict the noise added at each step, learning a time-conditional Fully Convolutional Network (U-Net) [28] *p*(*z_t_ −* 1*|z_t_*) that performs the reverse denoising process.

**Fig. 1.**
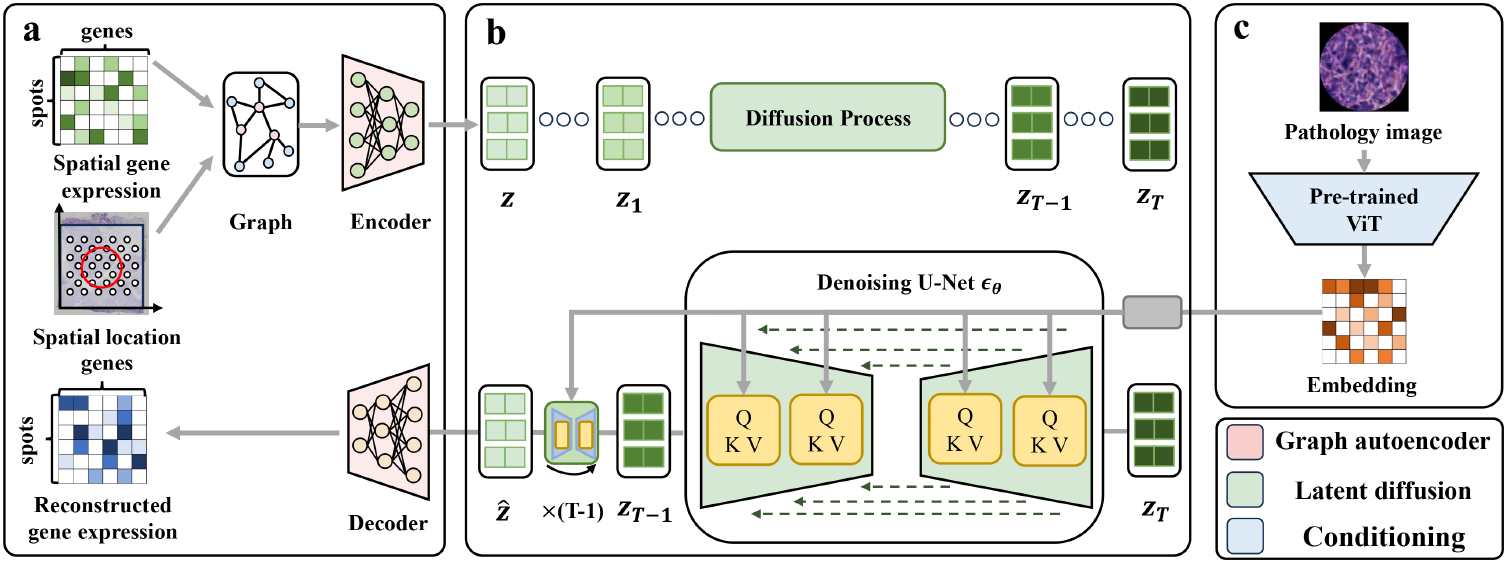
Overview of DiffuST. **a**, Spatial information compression and gene expression profiles reconstruction module. The module consists of a graph autoencoder that takes preprocessed spatial gene expression and a neighborhood graph constructed using spot coordinates as inputs. The encoder is utilized to compress the gene expression profiles and spatial information into a latent representation *z* and the decoder is utilized to reconstruct spatial gene expression from denoising latent representation *ẑ*. **b**, Latent representation denoising module. A latent diffusion model is designed to generate a denoising latent representation *ẑ*. The model consists of a diffusion process and a reverse process. During the diffusion process of DiffuST, Gaussian noise is iteratively added over *T* discrete steps until *z_T_* . In the reverse process, a time-conditional U-Net is trained to predict the noise added at each step. After training is complete, the diffusion model starts from *z_T_* and applies *T* steps of iterative denoising to generate a denoising latent representation *ẑ*. **c**, Image features conditioning mechanism module. The conditioning mechanism employs a pre-trained ViT model to obtain image features and utilize a cross-attention layer to incorporate them into the reverse diffusion process.

After training, to generate a denoising latent representation, the diffusion model initiates the process with *z_T_*, a highly noisy state of the original data. It then employs *T* steps of iterative denoising. During each step, the model uses the output from the previous denoising step as the input for the subsequent one. This iterative refinement continues until it reaches the final denoised representation *ẑ* (Fig 1b). Furthermore, to effectively use mapped imaging data to guide the denoising process, DiffuST employs a pre-trained ViT model to obtain image features and utilize a cross-attention layer to incorporate them into the reverse diffusion process (Fig 1c). Ultimately, the denoising latent representation *ẑ* is passed through the decoder of the graph autoencoder to obtain denoising spatial gene expression. Further details about the DiffuST can be found in the Methods section.

### 2.2 The denoising capability of DiffuST is more effective

Before evaluating the denoising capability of different methods, we first demonstrated the existence of extensive noise in spatial transcriptomics data. The details of the analysis are provided in the supplementary information.

After that, to evaluate the denoising capability of DiffuST, we conducted a benchmark analysis on the 10X Visium breast cancer dataset (Fig. 2a), comparing it with three state-of-art denoising methods (GraphST [19], MIST [20] and STACI [21]), which were published recently. We first used Moran’s I score [29] as the evaluation metric to assess whether the denoising data exhibit an organized spatial expression pattern. Moran’s I score close to one indicates a clear spatial pattern. The results indicated that DiffuST achieved a higher Moran’s I score compared to GraphST, MIST, and STACI (median of 0.885 for DiffuST against 0.792 for GraphST, 0.345 for MIST, and 0.815 for STACI), demonstrating DiffuST outperforms the other methods in spatial expression pattern recovery (Fig. 2b).

**Fig. 2.**
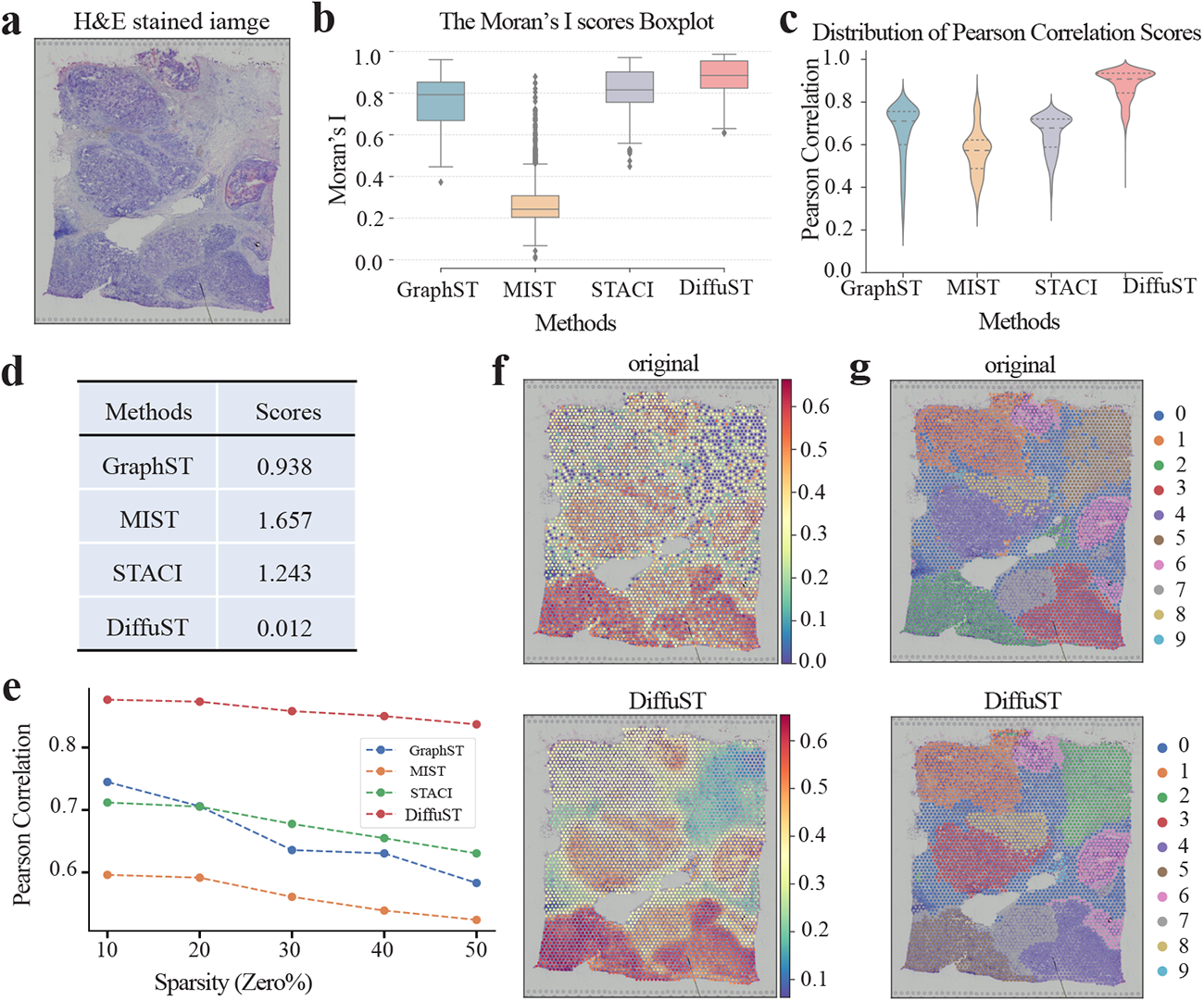
The denoising capability of DiffuST is more effective. **a**, The H&E stained image of the 10X Visium breast cancer dataset. **b**, Boxplot of Moran’s I score using DiffuST and other denoising methods. In the boxplot, the center line denotes the median, and box limits denote the upper and lower quartiles. **c**, Violin plot of the Pearson correlation between the original and denoising expression for the four methods. **d**, The table shows the average change of gene expression for each spot before and after the denoising process across the four methods. **e** The PCC score of each method with different dropouts. x-axis groups genes by different sparsity levels. **f**, Expression profile of gene HES4 before and after denoising generated by DiffuST. Expression profiles of gene HES4 after denoising by other methods are shown in Sup. Fig. 1. **g**, Clustering result before and after denoising generated by DiffuST. Clustering results before and after denoising generated by other methods are shown in Sup. Fig. 2.

To assess the impact of denoising on data consistency, we calculated the Pearson correlation coefficient (PCC) between the original and denoising expression [20]. Better denoising methods should have a higher PCC score. The result shows that DiffuST outperformed other methods in terms of data consistency while maintaining a clear spatial pattern (Fig. 2c). DiffuST exhibited an average PCC improvement of 21.3% compared with GraphST, 30.4% compared with MIST, and 22.2% compared with STACI. Additionally, to evaluate the stability of the denoising process, we compared the average change of gene expression for each spot before and after the denoising process (Fig. 2d). A better method is expected to have a lower value in this metric [30]. Among all compared methods, DiffuST exhibited minimal change before and after the denoising process, suggesting that it requires fewer changes in gene expression profiling to achieve optimal performance. Next, to investigate the capability of DiffuST against dropout noise, we assessed the performance of DiffuST and other methods on this dataset with different sparsity levels. To simulate dropout noise, a certain percentage of non-zero values in the expression matrix was set to zero. The Pearson correlation coefficient (PCC) was used to assess the performance of each method at different sparsity levels (Fig. 2e). Compared with other methods, DiffuST yielded stable PCC scores across different sparsity levels.

To intuitively show the denoising performance, we further visualized the expression profile of the breast cancer marker gene HES4 before and after denoising generated by different methods (Fig. 2f and Sup. Fig. 1) [31, 32]. DiffuST effectively amplifies the expression signals of HES4 in the tumor region, and the previously observed scattered expression pattern in the original expression profile disappears after denoising (Fig. 2g). While other denoising methods also strengthen the expression signals, the resulting profiles remain noisy and lack spatial continuity. Furthermore, we visualized the spot clustering result based on highly variable genes before and after denoising generated by different methods (Fig. 2g and Sup. Fig. 2). DiffuST accurately outlines complicated anatomical regions, while other methods tend to produce over-smoothed spatial domains that obscure fine-grained structures. Overall, all these results demonstrated the strength of DiffuST in denoising for spatial transcriptomics data.

### 2.3 Identifying tissue structures accurately with DiffuST

To evaluate the effectiveness of DiffuST in precisely identifying tissue structures, we applied it to a 10X Visium pancreatic cancer dataset initially annotated by Daniel et al [33]. The manual annotation identified regions of pancreatic intraepithelial neoplasia (PanIN), pancreatitis-like, and additional normal tissue (including normal duct)(Fig. 3a). We applied spatial clustering to the DiffuST-denoising data, selecting a configuration with k=3 clusters (Fig. 3b). This clustering result exhibited high congruence with the manually annotated tissue types. Notably, we find a central region of PanIN surrounded by the pancreatitis-like region, which is not accurately identified by other denoising methods (Fig. 3b and Sup. Fig. 3). This finding is in line with recent studies, which reported inflammation around the pancreatic ducts caused by PanIN [34, 35]. To quantify the accuracy of spatial clustering, we assessed the performance of spot clustering on both raw and denoising data generated by different methods, using the adjusted Rand index (ARI) and adjusted mutual information (AMI) as metrics [36]. DiffuST achieved the highest clustering accuracy compared to other methods, with an ARI of 0.636 and an AMI of 0.516, thereby demonstrating its superior ability to identify tissue structures (Fig. 3c).

**Fig. 3.**
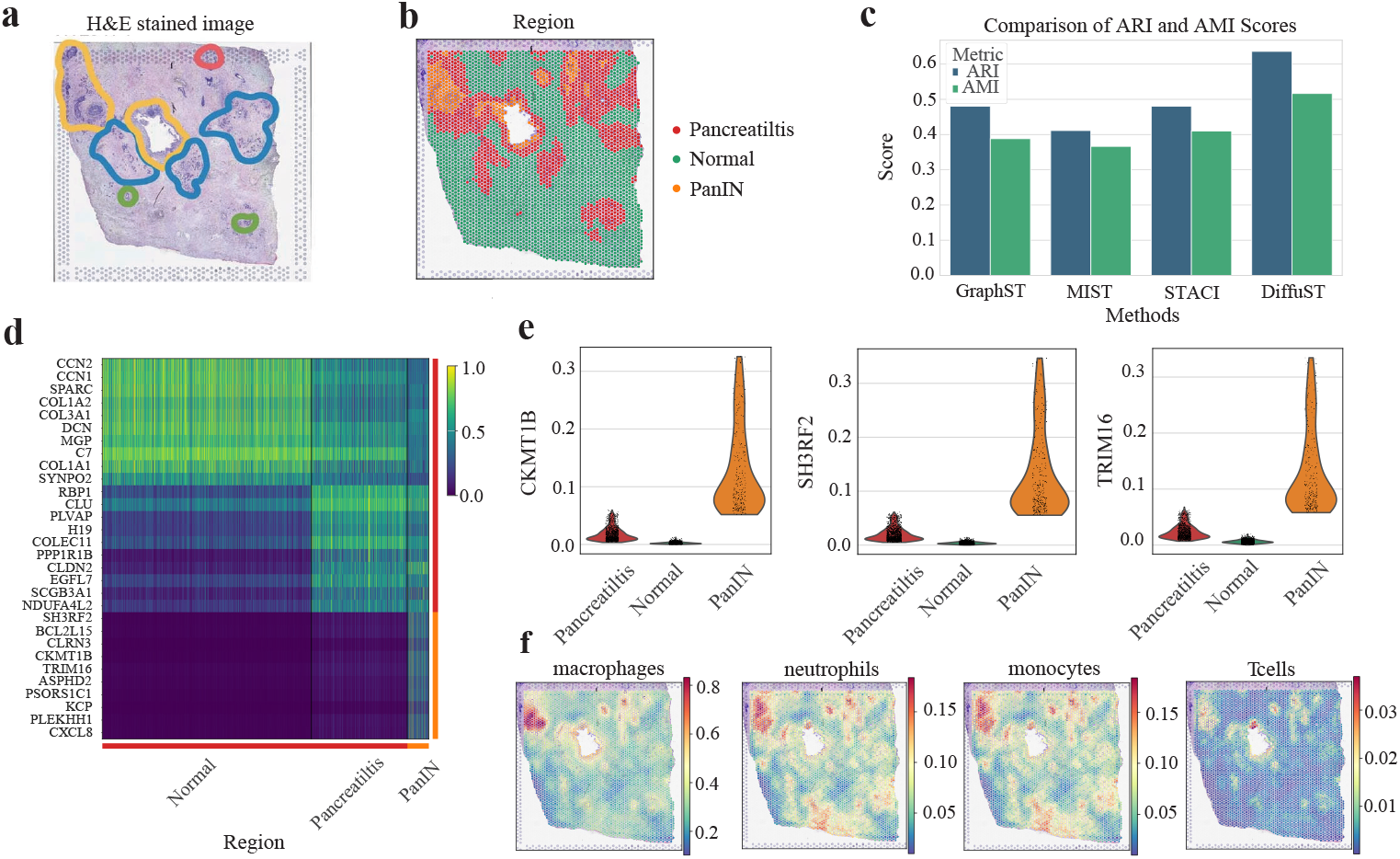
Identifying tissue structures accurately with DiffuST. **a**, Manual annotation of the 10X Visium pancreatic cancer dataset on a H&E stained image. The yellow region is PanIN, the blue region is pancreatitis-like and the other region is normal tissue (including normal duct). **b**, Visualization of the spatial domains detected by K-means spatial clustering on the denoising data by DiffuST. **c**, The comparison of ARI and AMI scores in the pancreatic cancer dataset for four methods. **d**, Differential expression analysis between the three tissue regions. The x-axis indicates different regions. **e**, The gene expression of CKMI1B, SH3RF2 and TRIM16 in different tissue regions. **f**, The spatial distributions of tumor, fibroblast, acinar, and T cells within the Diffusion-denoising gene expression profiles. The spatial distribution of different cell types on the original data and the denoising data by other methods are shown in Sup. Fig. 4.

Furthermore, we conducted a differential expression analysis between different tissue regions (Fig. 3d). The results revealed that PanIN regions exhibited distinct expression profiles, characterized by higher expression of genes associated with invasion and metastasis of pancreatic cancer such as SH3RF2, CKMT1B, and TRIM16 (Fig. 3e). Specifically, overexpression of SH3RF2 is associated with the invasion and metastasis of pancreatic cancer [37]. Overexpression of CKMT1B promotes tumor growth by inhibiting apoptosis in pancreatic cancer cells and downregulating the mitochondrial apoptotic pathway [38]. Additionally, TRIM16 enhances glycolysis to facilitate the migration of pancreatic cancer cells [39]. These findings provide strong support for the hypothesis that PanINs are a precursor to invasive pancreatic cancer at the aspect of gene expression [40, 41].

To evaluate whether the expression profiles denoised by DiffuST could provide a clearer spatial distribution of different cell types, we employed a signature-based strategy proposed by Rui Wu et al. to score the cell type enrichment in each spot based on marker genes from existing literature [33, 42]. This strategy utilized the average log-transformed normalization expression values of the genes as the corresponding cell type scores. Utilizing these cell type scores, we visualized the spatial distribution of cell types within the denoising gene expression profiles (Sup. Fig. 4). The denoising data revealed a more distinct spatial distribution pattern of cell types (Fig. 3f). Notably, we observed a strong molecular concordance between PanIN areas and acinar cells, consistent with existing studies suggesting PanIN develops from acinar cells that undergo acinar-ductal metaplasia [43, 44]. Moreover, compared to the pancreatitis regions, the PanIN regions present diverse immune cell types, including macrophages, neutrophils, monocytes, and plasma cells. This phenomenon indicates that PanIN lesions could induce an inflammatory response that promotes the development of pancreatic cancer [45]. These results collectively demonstrate that expression profiles denoised by DiffuST accurately identify tissue structures and provide a meaningful biological interpretation.

### 2.4 DiffuST facilitates the accurate inference of spatial cell-to-cell communications

Cell communication is the intricate process of mutual exchange and transmission of information between cells, facilitating their coordination and cooperation to maintain the normal functions of an organism [46]. Cell communication can manifest through various mechanisms, including direct cell-to-cell contact, secretion of signaling molecules between cells, and transmission of information through channels formed by intercellular connections [47]. To assess whether DiffuST denoising can enhance the spatial cell-cell interaction inference, we utilized it on a 10X Visium colorectal cancer dataset from recent studies (Fig. 4a) [48, 49]. Initially, the Leiden clustering method was applied to group the denoising spots, and these clusters were labeled as different regions, including fibroblast, smooth muscle, and tumor according to expert annotations (Fig. 4b).

Subsequently, cell communication analysis was conducted on both the original and DiffuST-denoising spatial expression data using CellChat to infer cellular communication between these three regions [46]. The cellular communication network revealed from data processed with DiffuST was more distinct than that from the original data, as indicated by significantly thicker edges (Fig. 4c and Sup. Fig. 5). This enhancement indicates that the denoising capability of DiffuST is helpful to uncover cell communication from noisy data. Detecting these signals accurately is crucial for understanding the complex intercellular interactions within colorectal cancer tissues [47]. Notably, despite the fibroblast region being close to both the tumor and smooth muscle regions, the communication signals between the fibroblast and the smooth muscle region were substantially weaker than those between the fibroblast and the tumor region. This difference emphasizes the effectiveness of the data denoised by DiffuST in enhancing intercellular communication signals and the accuracy of identifying intercellular interactions.

**Fig. 4.**
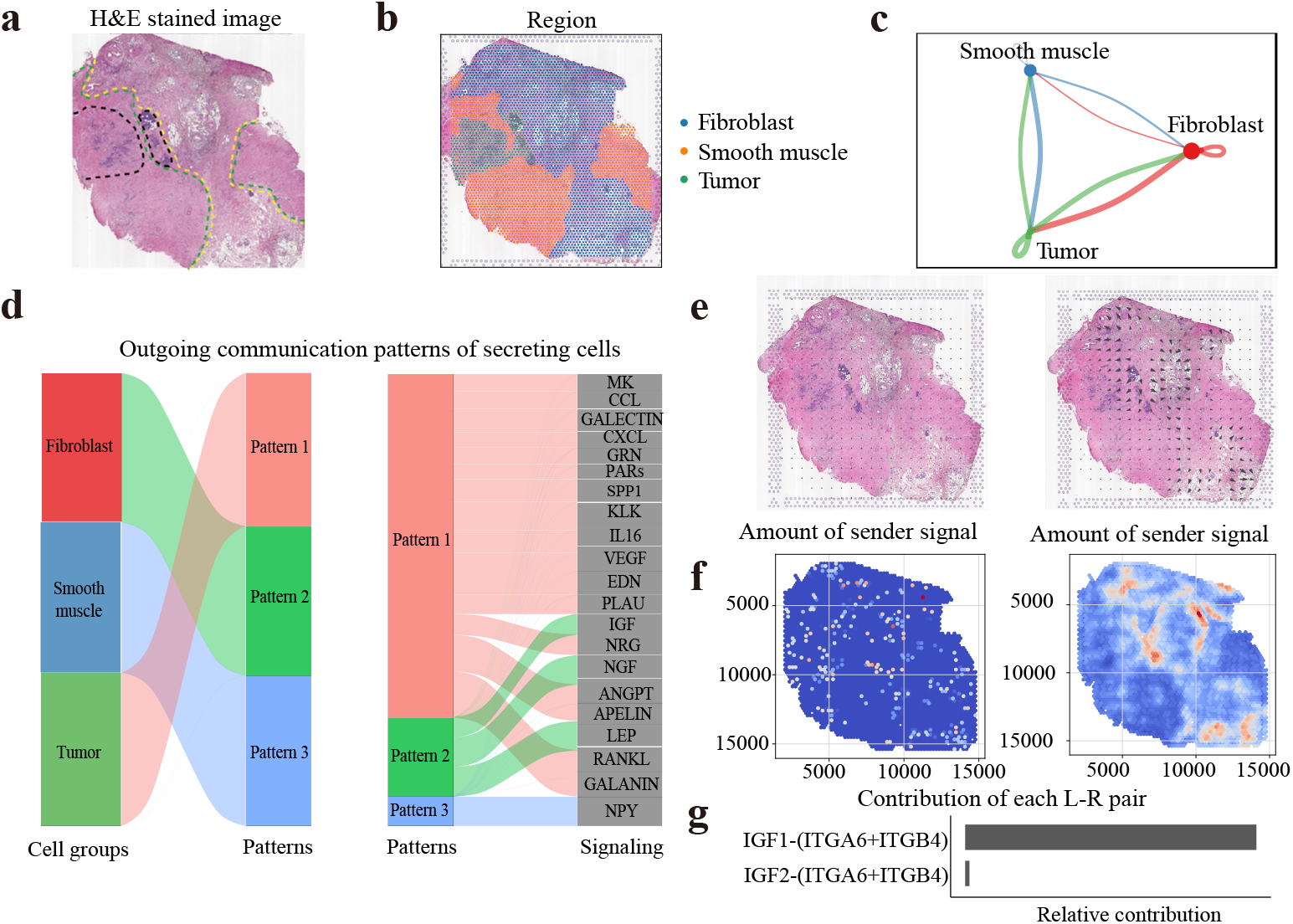
DiffuST facilitates the accurate inference of spatial cell-to-cell communications. **a**, Manual annotation of the 10X Visium colorectal cancer dataset on a H&E stained image. The yellow region is the fibroblast region, the green region is the smooth muscle region, and the black region is the tumor region. **b**, Visualization of the spatial domains detected by Leiden spatial clustering on the denoising data by DiffuST. **c**, The cellular communication network revealed from data processed with DiffuST. The cellular communication network revealed from the original data is shown in Sup. Fig. 5. The thickness of the line indicates the intensity of communication signals. Circle sizes are proportional to the number of cells in each cell group and edge width represents the number of communications. **d**, The visualization of the outgoing communication patterns from different regions. **e**, The visualization of the original and denoising direction of the IGF signaling pathways. The visualization of the original and denoising direction of the MK signaling pathways is shown in Sup. Fig. 6. **f**, The visualization of the original and denoising intensity of the IGF signaling pathways. The visualization of the original and denoising intensity of the MK signaling pathways is shown in Sup. Fig.7. **g**, The two IGF-related signaling pathways were distinctly recognized after DiffuST denoising.

To further explore the communication patterns between three distinct cell-type regions after denoising, we identified and visualized the outgoing communication patterns from these regions (Fig. 4d). The result revealed that the outgoing signaling from the tumor was characterized by Pattern 1, while that in fibroblast and smooth muscle was marked by Patterns 2 and 3, respectively. Fibroblasts, as the primary mesenchymal cells within colorectal cancer tissues, play a pivotal role in cancer promotion, cancer progression, and metastasis [50]. It was observed that Pattern 2 encompassed interactions involving insulin-like growth factor (IGF), neurotransmitter signals (NGF), and the interaction of neuroactive ligands with receptors (LEP). Specifically, the IGF signaling pathway plays a critical role in the development, survival, and progression of colorectal cancer [51]. Concurrently, evidence suggests that aberrant NGF expression in colorectal cancer may contribute to tumor growth and metastasis through neurotransmitter-mediated mechanisms [52]. Furthermore, a potential association exists between the LEP signaling pathway and colorectal cancer [53]. These insights underscore the efficacy of DiffuST in precisely inferring and elucidating the signaling pathways within different cellular landscapes.

To visually demonstrate the impact of DiffuST on enhancing inter-tissue signal transmission, we employed the COMMOT [47] to conduct a comparative analysis of direction and intensity in the IGF and MK signaling pathways before and after denoising (Fig. 4e, Fig. 4f, Sup. Fig. 6 and Sup. Fig. 7). The IGF signaling pathway was initiated in the fibroblast region, transmitting signals to the tumor region and smooth muscle region. Similarly, the MK pathway was traced from the tumor region targeting the smooth muscle region. This cannot be identified in the raw data. Furthermore, by comparing the number of ligand-receptor pairs within the signaling pathways before and after denoising, we confirmed a significant increase in the number of identified signals. For instance, no IGF-related signaling pathways were identified in the data before denoising. In contrast, two such pathways were distinctly recognized after DiffuST denoising (Fig. 4g).

### 2.5 Enhancing spatial transcriptomics deconvolution accuracy with DiffuST

To assess the efficacy of DiffuST in enhancing the accuracy of spatial transcriptomics deconvolution, we utilized it on a publicly available 10X Visium hepatocellular cancer (HCC) dataset [42] (Fig. 5a). The scRNA-seq data [54] employed for deconvolution consisted of approximately 74,000 cells classified into 11 distinct cell types (Fig. 5b). Furthermore, based on expert annotations, we used the Leiden clustering to delineate the spots into three primary tissue regions including the normal region, stromal region, and tumor region (Fig. 5c). We then used Tangram [14] to deconvolute the raw and DiffuST-denoising data using scRNA-seq as a reference. To intuitively show the impact of DiffuST on the accuracy of spatial transcriptomics deconvolution, we visualized the distribution of cell types within the gene expression profiles before and after denoising (Fig. 5d and Sup. Fig. 8). The denoising gene expression profiles accurately map different cell types to corresponding tissue regions. We observed a significantly higher distribution of Myeloid cells in tumor tissues than in normal and stromal tissues, which may be attributed to the vital role of Myeloid cells in supporting tumor initiation, progression, angiogenesis, and metastasis in hepatocellular cancer [55]. We also discovered a notable enrichment of Mast and Bi-Potent cells in the stromal regions, which were undetectable in the raw gene expression profiles. This suggests a potential involvement of these cell types in the formation and differentiation of fibroblasts within the stroma [56, 57]. Furthermore, we noted an enrichment of CD4+ T cells and CD8+ T cells in the tumor and normal tissues, consistent with the analysis performed by Rui et. al [58]. This demonstrates the effectiveness of DiffuST in improving the deconvolution accuracy of spatial transcriptomic data.

**Fig. 5.**
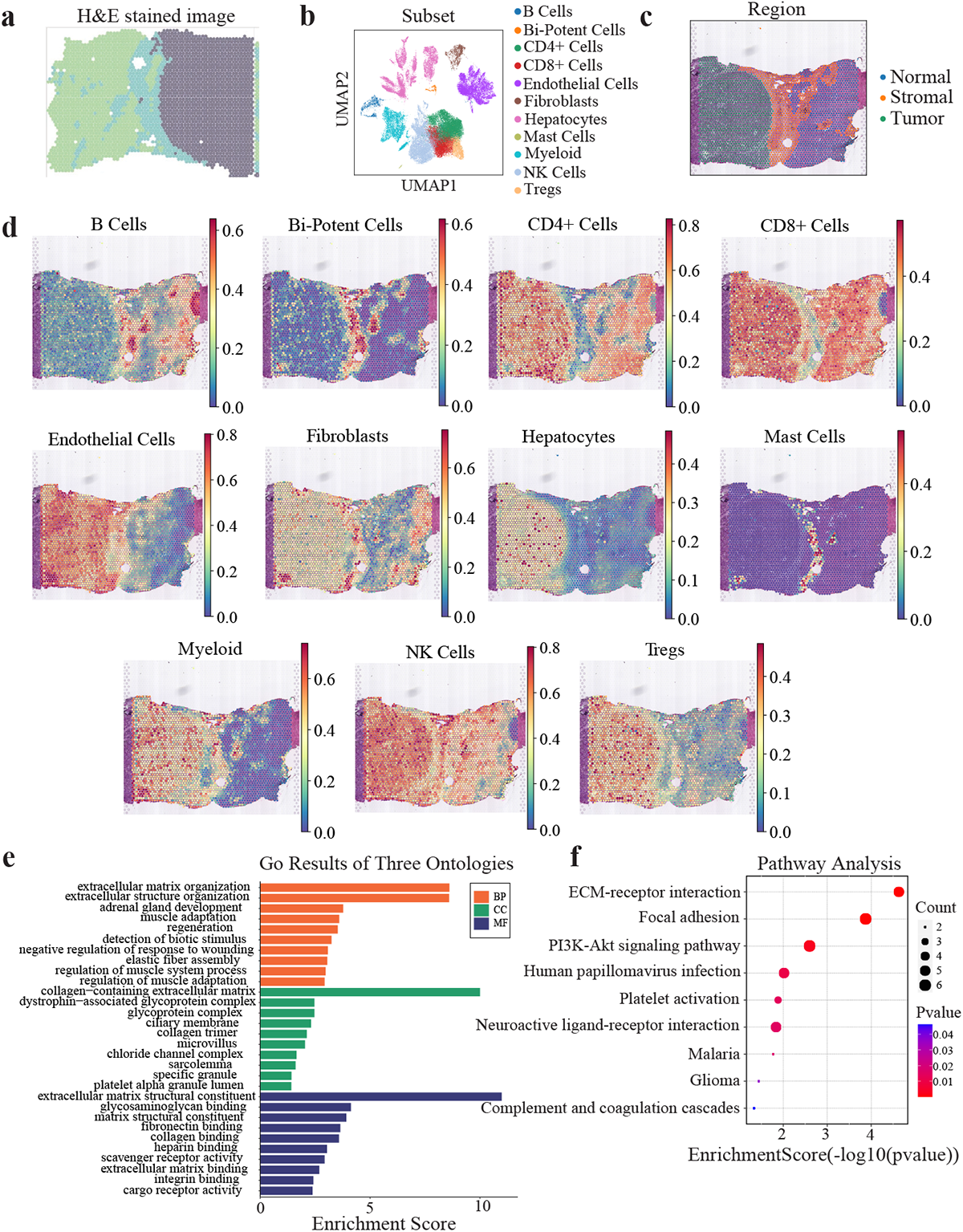
Enhancing spatial transcriptomics deconvolution accuracy with DiffuST. **a**, Manual annotation of the 10X Visium hepatocellular cancer dataset on a H&E stained image. The grey region is the tumor region, the blue region is the stromal region, and the green region is the normal region. **b**, Visualization of the UMAP of the scRNA-seq data employed for deconvolution. **c**, Visualization of the spatial domains detected by Leiden spatial clustering on the denoising data by DiffuST. **d**, The spatial distributions of major cell types within the gene expression profiles after denoising, namely B cells, Bi-Potent cells, CD4+ T cells, CD8+ T cells, Endothelial cells, Fibroblasts, Hepatocytes, Mast cells, Myeloid, NK cells, and Tregs. The spatial distributions of major cell types within the gene expression profiles before denoising are shown in Sup. Fig. 8. **e**, The results of the GO enrichment analysis within the stromal region. Statistical significance was assessed by the hypergeometric test, and p-values were adjusted by the Benjamini–Hochberg p-value correction algorithm. **f**, The results of the KEGG pathway analysis within the stromal region. The statistical significances were derived from hypergeometric tests and adjusted for multiple comparisons.

Subsequently, we explore key biological pathways and biological processes associated with tumor growth and progression in HCC [59]. We first identified differentially expressed genes, followed by conducting the Gene Ontology (GO) [60] enrichment analysis and Kyoto Encyclopedia of Genes and Genomes (KEGG) [61] pathway analysis on the stromal region. In the GO enrichment analysis, we noted that gene ontology terms related to the extracellular matrix (ECM) and its functions are dominant in the stromal region. This observation was consistent with existing research that indicates the stromal region consists of the ECM (FDR = 5 *×* 10*^−^*^5^, hypergeometric test adjusted by multiple comparisons) (Fig. 5e) [62]. Concurrently, there was a significant enrichment of gene ontology terms associated with glycosylation, such as glycoprotein complex formation, glycosaminoglycan binding, and matrix structural constituent glycosaminoglycan binding, within the same region. This result highlighted that glycosylation is associated with the development of HCC [63]. Furthermore, Wang et al. also demonstrated that dysregulated glycosylation regulates cancer growth, metastasis, stemness, immune evasion, and therapy resistance, and is regarded as a hallmark of HCC [64]. In KEGG pathway analysis, the significantly enriched pathways were the ECM-receptor interaction, focal adhesion, and PI3K-Akt signaling pathway (Fig. 5f). These enriched pathways have been confirmed to be consistent with HCC in existing research. Specifically, the activation of stromal cells and the excessive deposition of ECM leads to enhanced integrin signaling and stiffening of the liver tissue in cirrhosis and eventually the development of HCC [65]. Dysregulation of focal adhesion proteins often results in the acquired metastatic behavior of hepatocellular carcinoma [66]. Moreover, some evidence suggests that the activation of the PI3K/Akt signaling pathway may play a crucial role in enhancing the invasiveness and metastatic capability of HCC-resistant cells [67, 68]. These findings validate the efficacy of DiffuST in enhancing the accuracy of spatial transcriptomics deconvolution and explore the relationship between the results of enrichment analysis derived from denoising data and disease.

## 3 Discussion

Spatial transcriptomics is a technique that enables transcriptomics data to be acquired from intact tissue sections and provides spatial information. This technology addresses a key limitation of single-cell RNA sequencing (scRNA-seq), which is the absence of spatial resolution in the data [69]. It has revolutionized our comprehension of cellular heterogeneity, intercellular communication, and tissue structure by integrating spatial context with gene expression profiles [70]. Despite the considerable potential of spatial transcriptomics, the technique is hindered by existing extensive noise, which can impact downstream analyses and lead to erroneous interpretations [12, 71]. In this study, we developed a latent diffusion model named DiffuST to eliminate extensive noise in spatial transcriptomics by integrating gene expression profiles with spatial locations and corresponding imaging data. By learning a reverse diffusion process that transforms the noise distribution into the desired spatial gene expression distribution, DiffuST reduces noise in raw gene expression profiles and enhances the accuracy of downstream analysis from denoising spatial transcriptomics data.

While existing methods like SpaGE [15] and Tangram [14] achieve denoising by aggregating spatial gene expression profiles based on scRNAseq data, they may exhibit inconsistencies with the original spatial transcriptomics data in terms of expression values and correlations after denoising. Moreover, this type of method requires the matched (scRNA-seq) data from the same tissue, which needs extra experimental cost and is not always available. In contrast, DiffuST effectively integrates gene expression profiles, spatial information, and imaging features using the latent diffusion model, enabling denoising solely based on the spatial transcriptomics data itself. Although some existing methods like GraphST [19] and MIST [20] can utilize the location information and imaging data to enhance gene expression profiling, most of these methods primarily rely on positional information alone, limited in extracting global structural and semantic information even when utilizing imaging data. Unlike other methods that rely on direct pixel features from images, DiffuST employs a pre-trained ViT to effectively extract and integrate image features into the latent space, using a crossattention layer within the conditional denoising network to guide the denoising process. By testing on a variety of spatial transcriptomics datasets, we showed its superiority over existing denoising methods.

Importantly, our work suggests that denoising spatial transcriptomics data before any analysis will consistently improve the standard downstream analyses. By spatial clustering and differential expression analysis on pancreatic cancer data, we unveiled the spatial domain of PanIN and its potential associations with the invasive functions of pancreatic cancer. Furthermore, by comparing the transmission of intercellular interaction signals before and after denoising, we discovered that spatial transcriptomics noise significantly impacts the accurate inference of intercellular communication signals. Finally, through a series of deconvolution experiments, we uncovered immune heterogeneity and potential regulatory pathways between distinct regions in hepatocellular carcinoma. Overall, DiffuST is a powerful method for spatial transcriptomics denoising, which enables enhanced downstream analysis and provides valuable insights into complex biological systems.

## 4 Conclusion

Spatial transcriptomics technologies have enabled comprehensive measurements of gene expression profiles while retaining spatial information and matched pathology images. Current denoising methods often emphasize spatial localization to enhance gene expression data, ignoring the rich structural and semantic information present in corresponding histology images. Here, we develop a latent diffusion model DiffuST to denoise spatial transcriptomics. By testing on various spatial transcriptomics datasets, we showed its superiority over existing denoising methods. The results demonstrated that denoising spatial transcriptomics data consistently improves the standard downstream analyses, including tissue structure identification, inference of cell-to-cell communications, and deconvolution of cell types. Overall, DiffuST is a powerful method for spatial transcriptomics denoising, which enables enhanced downstream analysis and provides valuable insights into complex biological systems.

## 5 Availability of data and materials

All datasets used in this work are publicly available from the following sources: The 10X Visium prostate cancer dataset and the 10X Visium breast cancer dataset were obtained from https://www.10xgenomics.com/datasets. The 10X Visium pancreatic cancer dataset was obtained from https://data.humantumoratlas.org/. The 10X Visium colorectal cancer dataset was available from http://www.cancerdiversity.asia/scCRLM/. The 10X Visium hepatocellular cancer dataset was available from https://ngdc.cncb.ac.cn/gsa-human/browse/HRA000437 and the reference scRNA-seq data used for spatial transcriptomics deconvolution was available from https://data.mendeley.com/datasets/6wmzcskt6k/1.

## 7 Author contributions

Conceptualization, J.P.; methodology, J.P. and S.J.; experimentation, S.J.; writing—original draft, S.J.; writing—review and editing, J.P., S.J., Y.D., D.L., X.Z., T.W., and Y.W.; supervision, J.P. and Y.D.

