## supplementary for "DiffuST: a latent diffusion model for spatial transcriptomics denoising"

### 1 Spatial transcriptomics data exists extensive noise

We initiated our analysis by assessing noise in the spatial transcriptomics using a 10X Visium prostate cancer dataset, along with matched Immunofluorescence (IF) images stained with Iba1, Vimentin, and DAPI [1]. When comparing IF staining images and the Iba1 antibody expression intensity with the gene expression profiling of AIF1 (the gene that encodes Iba1) derived from transcriptome sequencing (as shown in Sup. Fig. 9a), there is a poor correlation between the protein expression levels of Iba1 and the RNA expression levels of AIF1. This suggests that noise exists in gene expression profiling. There are lots of zero-values for the AIF1 RNA expression, clearly showing high levels of dropouts in AIF1 expression (Sup. Fig. 9b). This suggests that one source of noise in ST is dropouts, similar to the scRNA-seq [2, 3]. Additionally, we investigated the impact of different sequencing qualities on the dropout rate. The total sequencing depth of each spot can be used to represent the sequencing quality of a spot. We divided all spots into four categories based on the total sequencing depths and plotted the gene expression distributions of ESR1 for each category. Under different categories, we expected the gene expression of spots to follow a similar distribution. However, there is a dramatic difference between gene expression distributions with different categories (Sup. Fig. 9c). The percentages of zero counts increased with the decrease in the sequencing quality. After denoising by different models, the percentages of zero counts from different sequencing qualities were significantly reduced, and the gene expression distributions of spots became consistent (Sup. Fig. 10). These results suggest that denoising can eliminate the impact of different sequencing qualities.

To demonstrate the noise in spatial transcriptomics data may also arise from high gene expression inflation [4], we analyzed another 10X Visium human breast cancer dataset. In breast tissue, multiple island-like structures exist (darker areas in the H&E-stained tissue images), which represent luminal cell compartments surrounded by connective tissue [5, 6] (Sup. Fig. 9d and Sup. Fig. 11). However, the expression of ESR1, a marker gene for luminal cells, in this Visium dataset only shows a weak correlation with the island-like structures on H&E-stained tissue sides. The weak correlation observed cannot be entirely attributed to dropouts, considering that the expression level of the gene ESR1 remains elevated in some regions, like the connective tissue surrounding the luminal cell compartment, where it is not typically expected.

Finally, we verified that the noise within ST datasets causes inaccurate estimation of gene-gene spatial correlation, which is the fundamental element in many analyses [7, 8]. To assess the impact of noise on spatial co-expression patterns, we take the Pearson correlation coefficient between a pair of genes, MKI67 and CCNB1, as an example. They have been confirmed to exhibit high co-expression at the single-cell level in breast cancer [9]. However, the original spatial transcriptomics data yielded a correlation score of only 0.16, which was lower than expected (Sup. Fig. 9e). For comparison, we applied different denoising methods to this data. The results showed a significant enhancement in correlation scores, consistent with findings from another single-cell study [10]. This case demonstrates that co-expression patterns are not significant in the original ST data due to the presence of noise (Sup. Fig. 9e and Sup. Fig. 12).

#### 2 Supplementary Figures

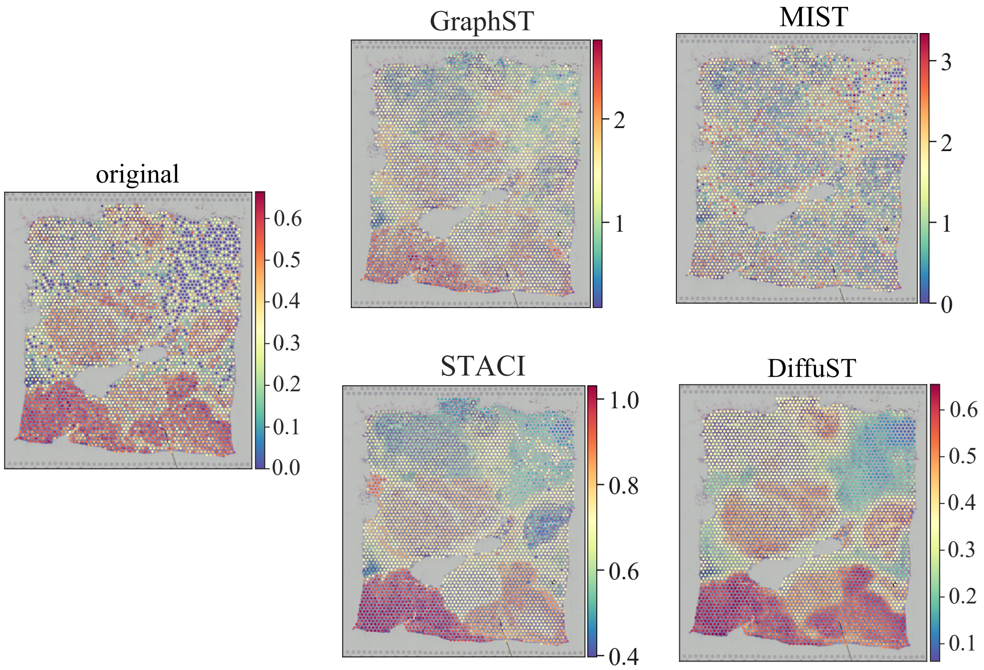


**Supplementary Figure1.** Expression profiles of gene HES4 before and after denoising generated by GraphST, STACI, MIST, and DiffuST.

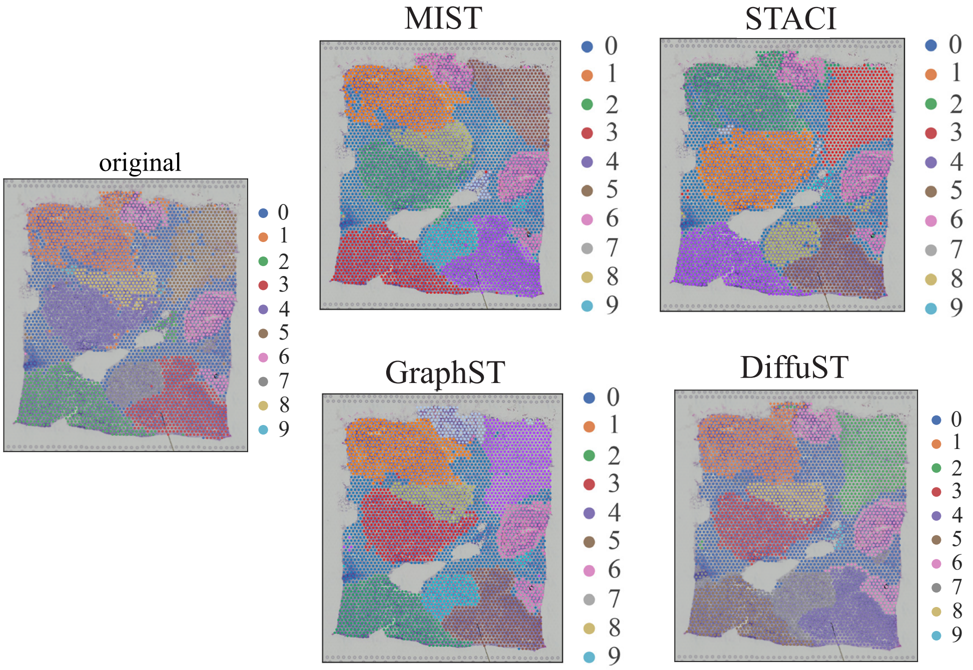


**Supplementary Figure2.** Clustering results before and after denoising generated by GraphST, STACI, MIST, and DiffuST.

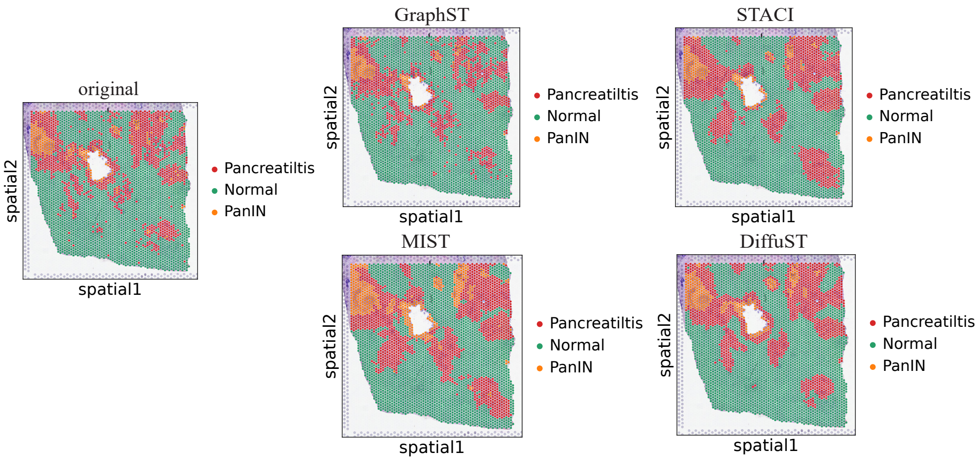


**Supplementary Figure3.** Visualization of the spatial domains detected by K-means spatial clustering on the denoising data by GraphST, STACI, MIST, and DiffuST.

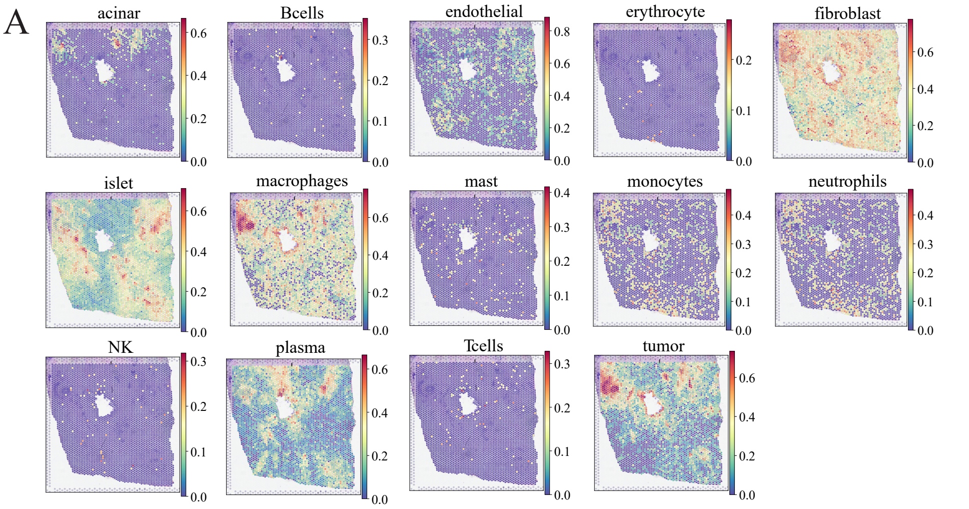


**Supplementary Figure4A.** The spatial distribution of acinar, B-cells endothelial, erythrocyte, fibroblast, islet, macrophages, mast, monocytes, neutrophils, NK, plasma, T-cells, and tumor within the original gene expression profiles.

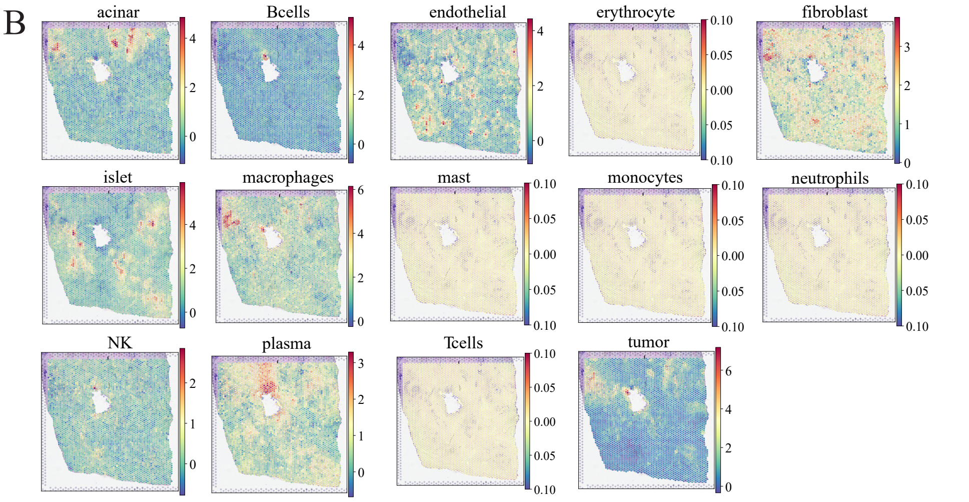


**Supplementary Figure4B.** The spatial distribution of acinar, B-cells endothelial, erythrocyte, fibroblast, islet, macrophages, mast, monocytes, neutrophils, NK, plasma, T-cells, and tumor within the GraphST-denoising gene expression profiles.

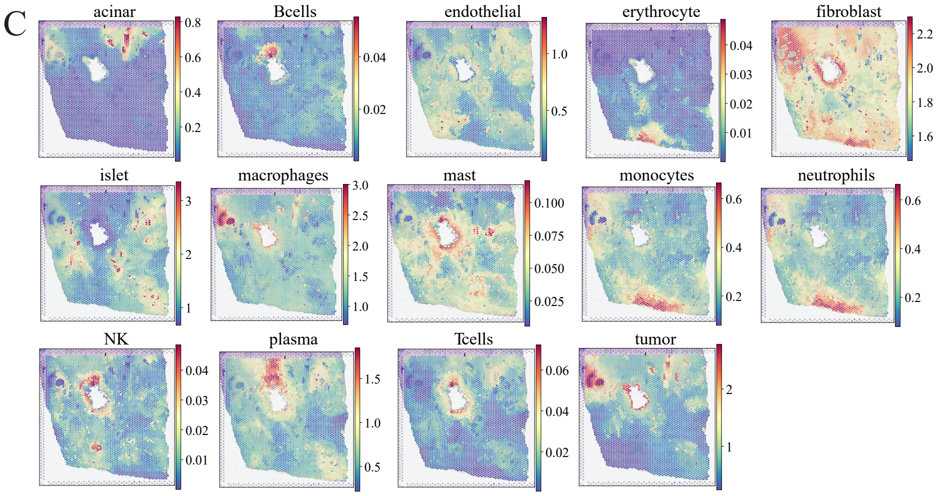


**Supplementary Figure4C.** The spatial distribution of acinar, B-cells endothelial, erythrocyte, fibroblast, islet, macrophages, mast, monocytes, neutrophils, NK, plasma, T-cells, and tumor within the STACI-denoising gene expression profiles.

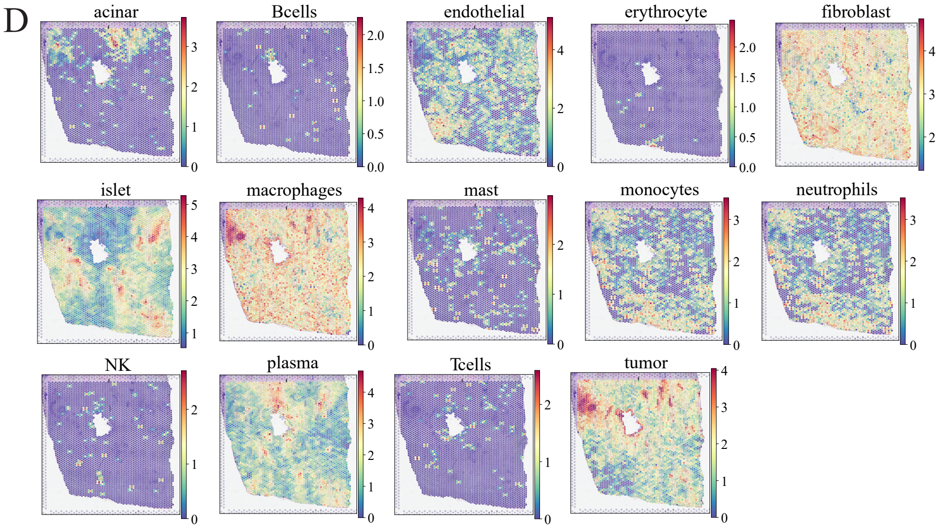


**Supplementary Figure4D.** The spatial distribution of acinar, B-cells endothelial, erythrocyte, fibroblast, islet, macrophages, mast, monocytes, neutrophils, NK, plasma, T-cells, and tumor within the MIST-denoising gene expression profiles.

E

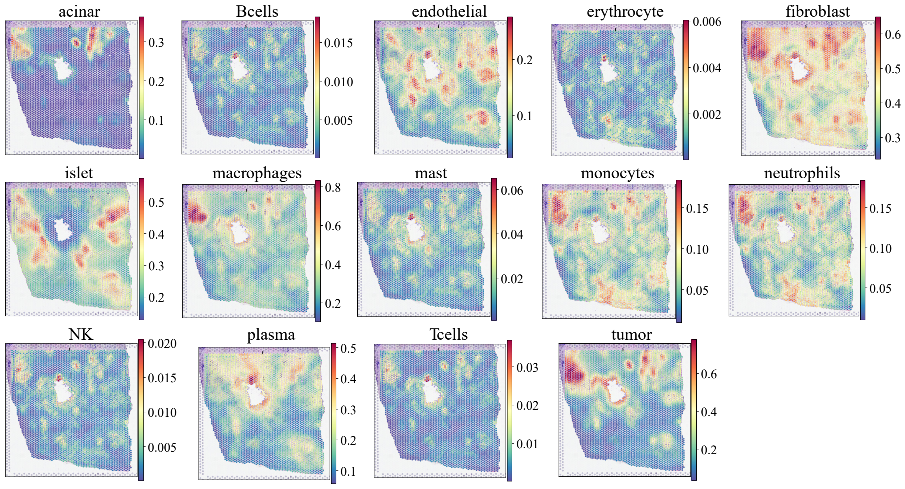


**Supplementary Figure7E.** The spatial distribution of acinar, B-cells endothelial, erythrocyte, fibroblast, islet, macrophages, mast, monocytes, neutrophils, NK, plasma, T-cells, and tumor within the Diffusion-denoising gene expression profiles.

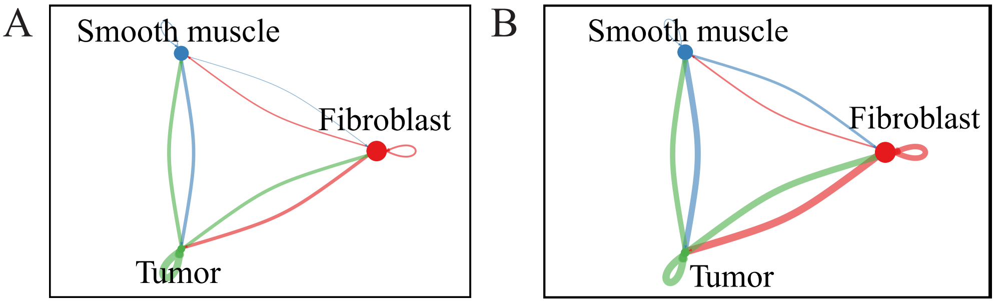


**Supplementary Figure5. A**. The cellular communication network is revealed from the original data. **B**. The cellular communication network revealed from data processed with DiffuST.

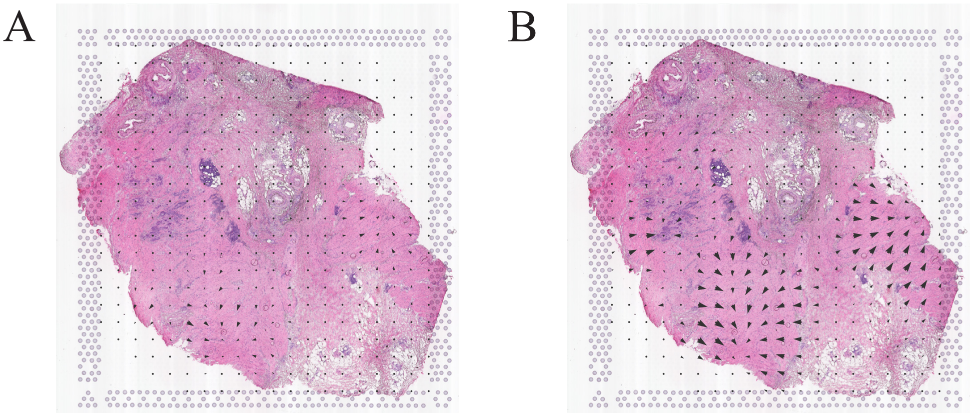


**Supplementary Figure6. A**. The visualization of the original direction of the MK signaling pathways. **B**. The visualization of the denoising direction of the MK signaling pathways.

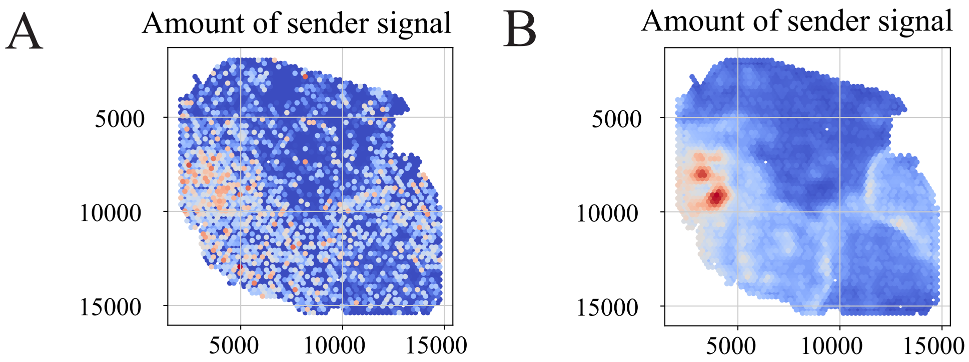


**Supplementary Figure7. A**. The visualization of the original intensity of the MK signaling pathways. **B**. The visualization of the denoising intensity of the MK signaling pathways.

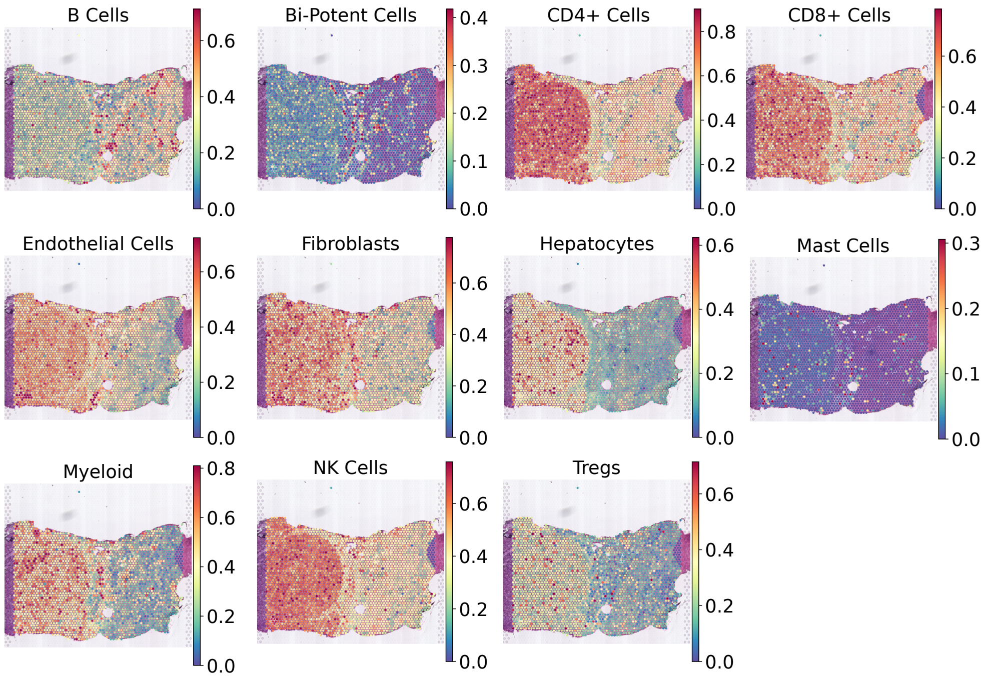


**Supplementary Figure8.** The spatial distributions of major cell types within the gene expression profiles before denoising, namely B cells, Bi-Potent cells, CD4+ T cells, CD8+ T cells, Endothelial cells, Fibroblasts, Hepatocytes, Mast cells, Myeloid, NK cells, and Tregs.

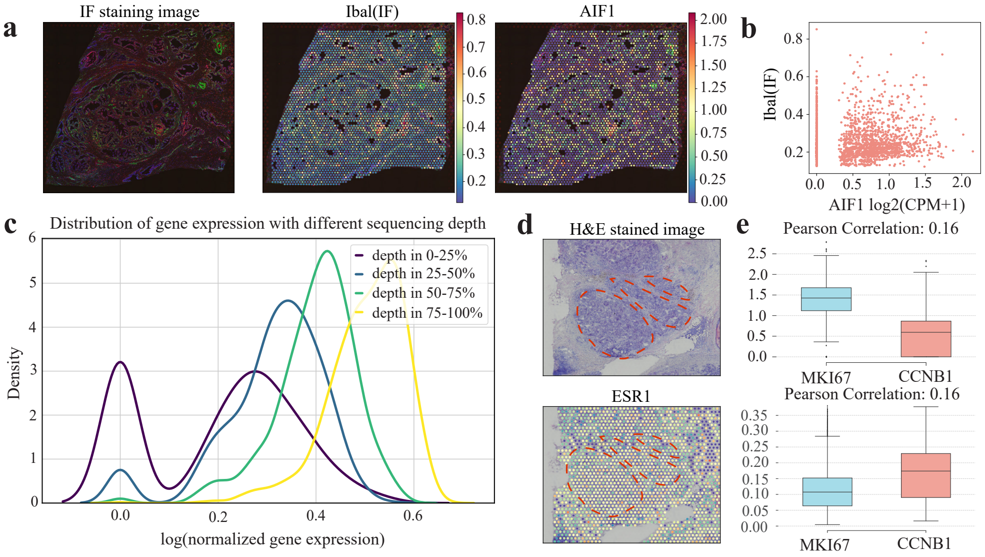


**Supplementary Figure9. Spatial transcriptomics data exists extensive noise.** **a**, IF staining image, the Iba1 antibody expression intensity, and the RNA expression levels of AIF1 of the 10X Visium prostate cancer dataset. **b**, Scatterplots showing the correlation between AIF1 RNA expression and the intensity of Iba antibody expression. The x-axis shows the AIF1 RNA expression level (unit = log2(CPM+1)). **c**, The gene expression distributions of ESR1 under different sequencing depths. The y-axis shows the gene expression density of ESR1. **d**, Example island-like structure with a poor agreement with ESR1 expression from the 10X Visium human breast cancer. The red circle marks the luminal cell compartment. **e**, Boxplot of Pearson correlation coefficient between MKI67 and CCNB1 expression from the original and DiffuST-denoising data.

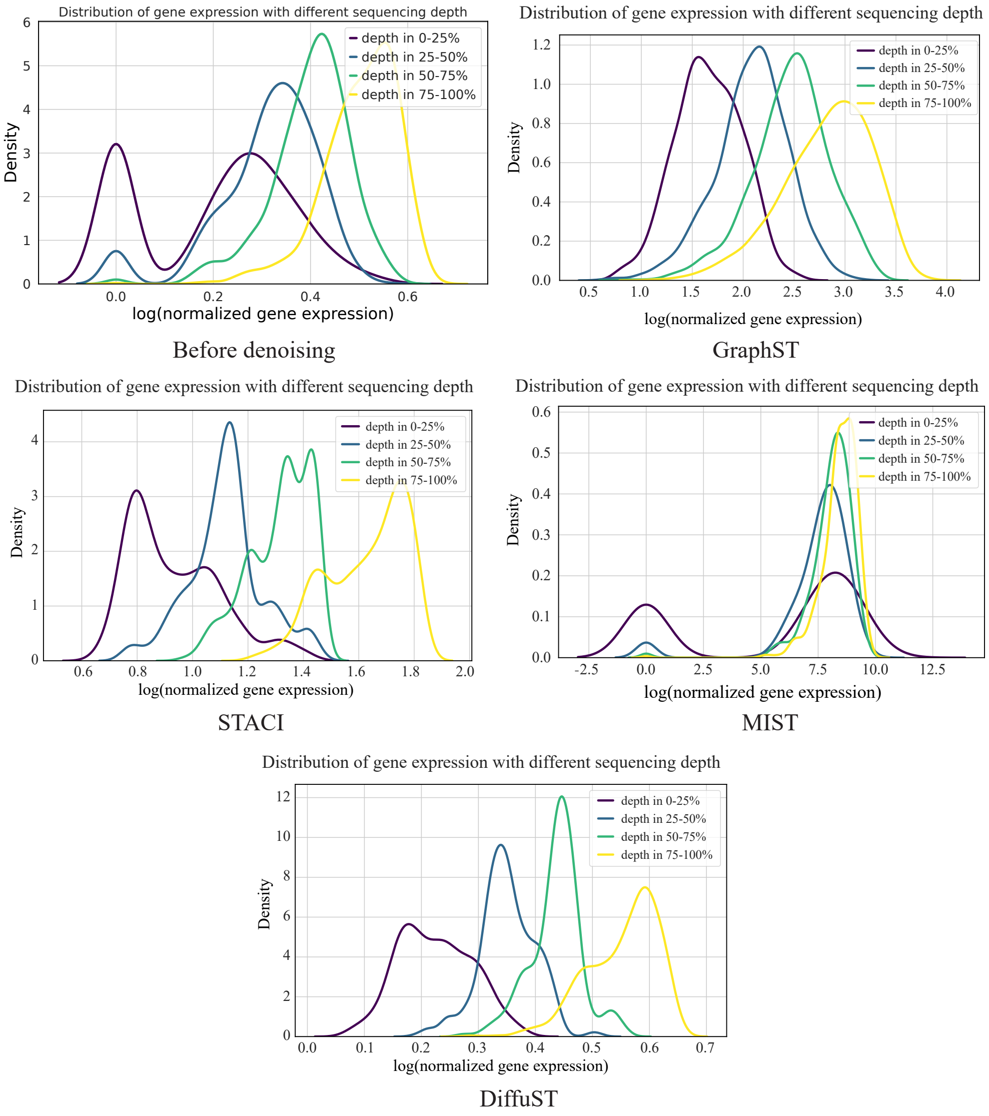


**Supplementary Figure10.** The gene expression distributions of ESR1 under different sequencing depths before and after denoised by GraphST, STACI, MIST, and DiffuST. The y-axis shows the gene expression density of ESR1.

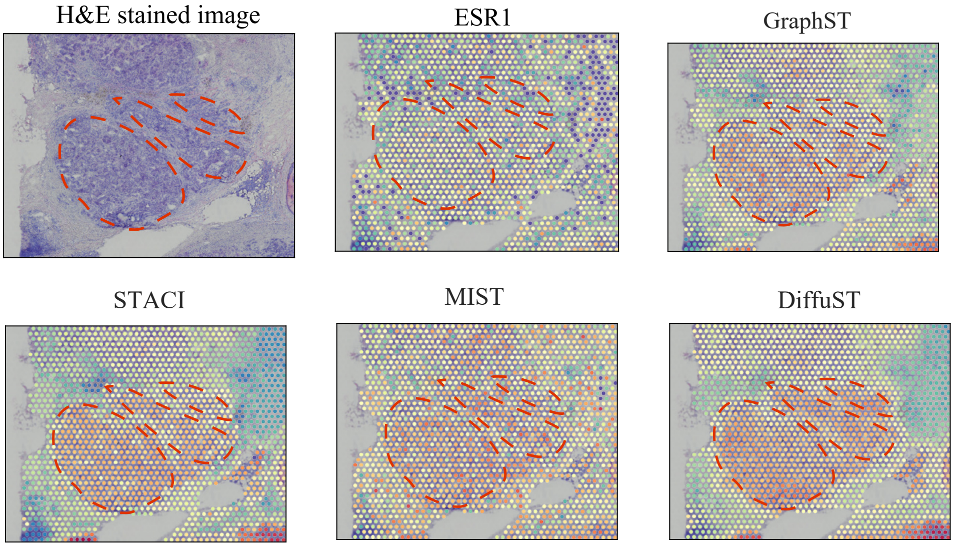


**Supplementary Figure11.** Comparison of the island-like structure before and after denoised by GraphST, STACI, MIST, and DiffuST. The first figure is an H&E-stained image of the island-like structure from the 10X Visium breast cancer dataset. The other figures are the island-like structure before and after denoised by GraphST, STACI, MIST, and DiffuST. The red circle marks the luminal cell compartment.

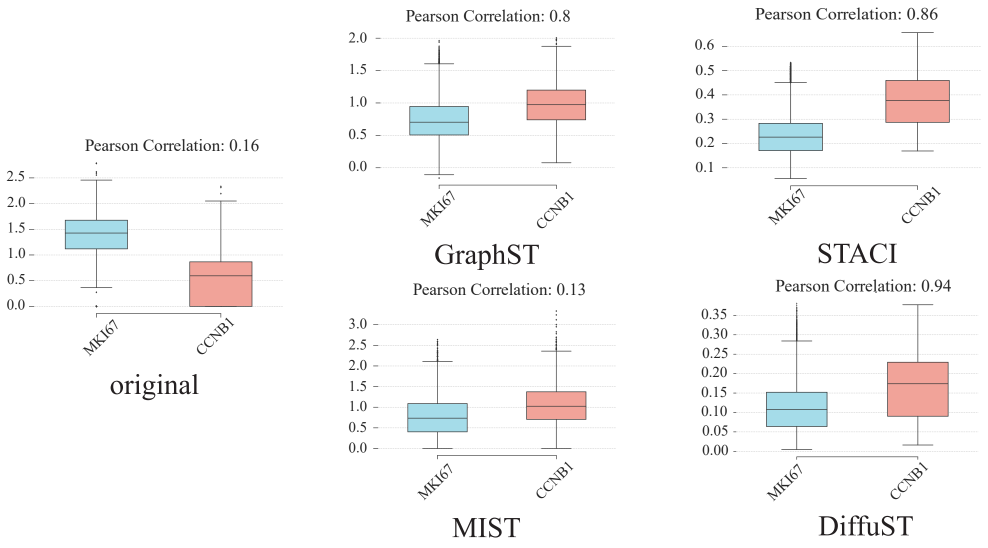


**Supplementary Figure12.** Boxplots of the Pearson correlation coefficient between MKI67 and CCNB1 expression from the data denoised by GraphST, STACI, MIST, and DiffuST.

### 3 Supplementary Tables

**Supplementary Table 1**: Description of all ST and SC datasets used in this study

| Dataset | Number of spots/cells | Number of genes | Sparsity levels |
|---|---|---|---|
| Prostate cancer ST | 3460 | 17943 | 26.57% |
| Breast cancer ST | 3798 | 36601 | 15.36% |
| Pancreatic cancer ST | 4992 | 36601 | 7.85% |
| Colorectal cancer ST | 4007 | 36601 | 5.55% |
| Hepatocellular cancer ST | 2791 | 33538 | 13.06% |
| Hepatocellular cancer SC | 73589 | 2608 | 100% |

This table provides a clear overview of the datasets used in the study, their type (ST or SC), the number of spots/cells, the number of genes, and the sparsity level, which indicates the proportion of zero-expression values in each dataset.

**Supplementary Table 2**: The structure of the graph autoencoder in DiffuST

| Component | Stage | Layer detail | Input size | Output size |
|---|---|---|---|---|
| Encoder | Compress features | GCNConv + Leaky ReLU | 3000 | 512 |
| | Calculate *mu* | GCNConv + Leaky ReLU | 512 | 128 |
| | Calculate *logvar* | GCNConv + Leaky ReLU | 512 | 128 |
| Decoder | Decompress features | GCNConv + Leaky ReLU | 128 | 512 |
| | Calculate *pi* | GCNConv + Sigmoid | 512 | 3000 |
| | Calculate *theta* | GCNConv + Softplus | 512 | 3000 |
| | Calculate *mean* | GCNConv + Exp | 512 | 3000 |

This table provides a structured overview of the layers and their configurations for each stage of the graph autoencoder in DiffuST.

**Supplementary Table 3**: The description and setting of the hyperparameters for latent diffusion model

| Hyperparameter | Default | Description |
|---|---|---|
| Input size | 128 | The dimension of the input |
| timesteps | 1000 | The number of steps used to add noise |
| DDIM steps | 50 | The number of denoising steps |
| Strength | 0.6 | The strength of the noise used in the diffusion process |
| Batch Size | 64 | The number of training samples |
| Iters | 300 | The epochs of the training |
| Learning Rate | 1e-4 | Learning rate of the model |
| Schedule | DDIM | A scheduler to be used in combination with U-Net to denoise the encoded features |
